# Refactoring the formicamycin biosynthetic gene cluster to make high-level producing strains and new molecules

**DOI:** 10.1101/2020.09.15.275776

**Authors:** Rebecca Devine, Hannah McDonald, Zhiwei Qin, Corinne Arnold, Katie Noble, Govind Chandra, Barrie Wilkinson, Matthew I. Hutchings

**Affiliations:** Department of Molecular Microbiology, John Innes Centre, Norwich Research Park, Norwich, UK, NR4 7UH

## Abstract

The formicamycins are promising antibiotics with potent activity against Gram-positive pathogens including VRE and MRSA and display a high barrier to selection of resistant isolates. They were first identified in *Streptomyces formicae* KY5, which produces the formicamycins at low levels on solid agar but not in liquid culture, thus hindering further investigation of these promising antibacterial compounds. We hypothesised that by understanding the organisation and regulation of the *for* biosynthetic gene cluster, we could rationally refactor the cluster to increase production levels. Here we report that the *for* biosynthetic gene cluster consists of 24 genes expressed on nine transcripts. Seven of these transcripts, including those containing all the major biosynthetic genes, are repressed by the MarR-regulator ForJ which also controls the expression of the ForGF two-component system that initiates biosynthesis. A third cluster-situated regulator, ForZ, autoregulates and controls production of the putative MFS transporter ForAA. Consistent with these findings, deletion of *forJ* increased formicamycin biosynthesis 5-fold, while over-expression of *forGF* in the Δ*forJ* background increased production 10-fold compared to the wild-type. De-repression by deleting *forJ* also switched on biosynthesis in liquid-culture and induced the production of two novel formicamycin congeners. By combining mutations in regulatory and biosynthetic genes, six new biosynthetic precursors with antibacterial activity were also isolated. This work demonstrates the power of synthetic biology for the rational redesign of antibiotic biosynthetic gene clusters both to engineer strains suitable for fermentation in large scale bioreactors and to generate new molecules.

**Importance:** Antimicrobial resistance is a growing threat as existing antibiotics become increasingly ineffective against drug resistant pathogens. Here we determine the transcriptional organisation and regulation of the gene cluster encoding biosynthesis of the formicamycins, promising new antibiotics with activity against drug resistant bacteria. By exploiting this knowledge, we construct stable mutant strains which over-produce these molecules in both liquid and solid culture whilst also making some new compound variants. This will facilitate large scale purification of these molecules for further study including in vivo experiments and the elucidation of their mechanism of action. Our work demonstrates that understanding the regulation of natural product biosynthetic pathways can enable rational improvement of the producing strains.

## Introduction

Almost half of all known antibiotics are derived from the specialised metabolites of filamentous actinomycetes, particularly *Streptomyces* species, many of which were discovered during the Golden Age of antibiotic discovery that occurred between 1940 and 1960 ^1^. Since this time, few new classes of antibiotics have been introduced into the clinic and increasing problems of antimicrobial resistance pose a significant threat to modern medicine. Many synthetic antibiotics have failed to progress through clinical trials, so interest has returned to natural products ^2^. *Streptomyces* are primarily known as soil bacteria, however, their ability to produce antibiotics makes them competitive in a wide range of environments. By searching under-explored environments and ecosystems new species can be isolated that produce new specialised metabolites ^3^. Furthermore, advances in genomic techniques has revealed that bacterial and fungal species encode many more biosynthetic gene clusters (BGCs) than previously thought, with only around 10% being expressed under laboratory conditions, meaning many more specialised metabolites remain to be discovered from their cryptic BGCs. Isolating new species from under-explored environmental niches and mining their genomes for novel BGCs is therefore a promising route towards finding new molecules for antibiotic development ^4^. Following this hypothesis, we previously isolated a number of actinomycetes from the domatia of the African fungus-growing plant-ant *Tetraponera penzigi,* including the new species *Streptomyces formicae* KY5 ^5^. The genome of *S. formicae* encodes at least 45 specialised metabolite BGCs including a Type 2 polyketide synthase (PKS) BGC that is responsible for the biosynthesis of formicamycins ^6^. These antibiotics are potent inhibitors of vancomycin resistant enterococci (VRE) and MRSA, with no resistance observed *in vitro* ^7^.

Previous work on the formicamycin (*for*) BGC has shown that the encoded biosynthetic pathway makes two distinct families of compounds in addition to the formicamycins. The fasamycins are biosynthetic precursors of the formicamycins that also exhibit antibacterial activity; they have been isolated from a number of actinomycete strains and given the alternative names accramycins, naphthacemycins and streptovertimycins ^8–11^. In addition to the fasamycins, the formicapyridines are shunt metabolites produced when the cyclisation stage of the biosynthetic pathway is derailed ^7,12^. Conversion of fasamycin precursors into formicamycins involves a unique two-step ring-expansion, ring-contraction pathway that proceeds via a Baeyer-Villigerase derived lactone intermediate which undergoes a unique reduction Favorskii-like rearrangement ^13^ (**Figure 1**). In this work, we aimed to understand how *S. formicae* KY5 regulates the production of formicamycins with the view to refactoring the BGC to produce increased titres of these potentially valuable antibiotics. We show that the *for* BGC consists of 24 genes expressed on nine transcripts and is controlled by the combined actions of three cluster-situated regulators (CSRs). The MarR family transcriptional regulator ForJ represses the expression of the majority of the biosynthetic genes, while the two-component system (TCS) ForGF is required to activate formicamycin biosynthesis. A third CSR, the MarR family regulator ForZ, appears to auto-repress its own expression and activate expression of the putative, divergent MFS transporter gene *forAA*. Deletion of the *forGF* operon abolished the production of fasamycins and formicamycins in the wild-type strain while deleting *forJ* increased formicamycin titres approximately 5-fold. Introducing a second copy of *forGF* into the ΔforJ mutant increased production of formicamycins to approximately 10 times the wild-type levels. De-repression of the BGC by deleting *forJ* also induced the production of fasamycins and formicamycins in liquid culture, as well as inducing the production of two novel formicamycin congeners and six new precursor fasamycins.

**Figure 1.**
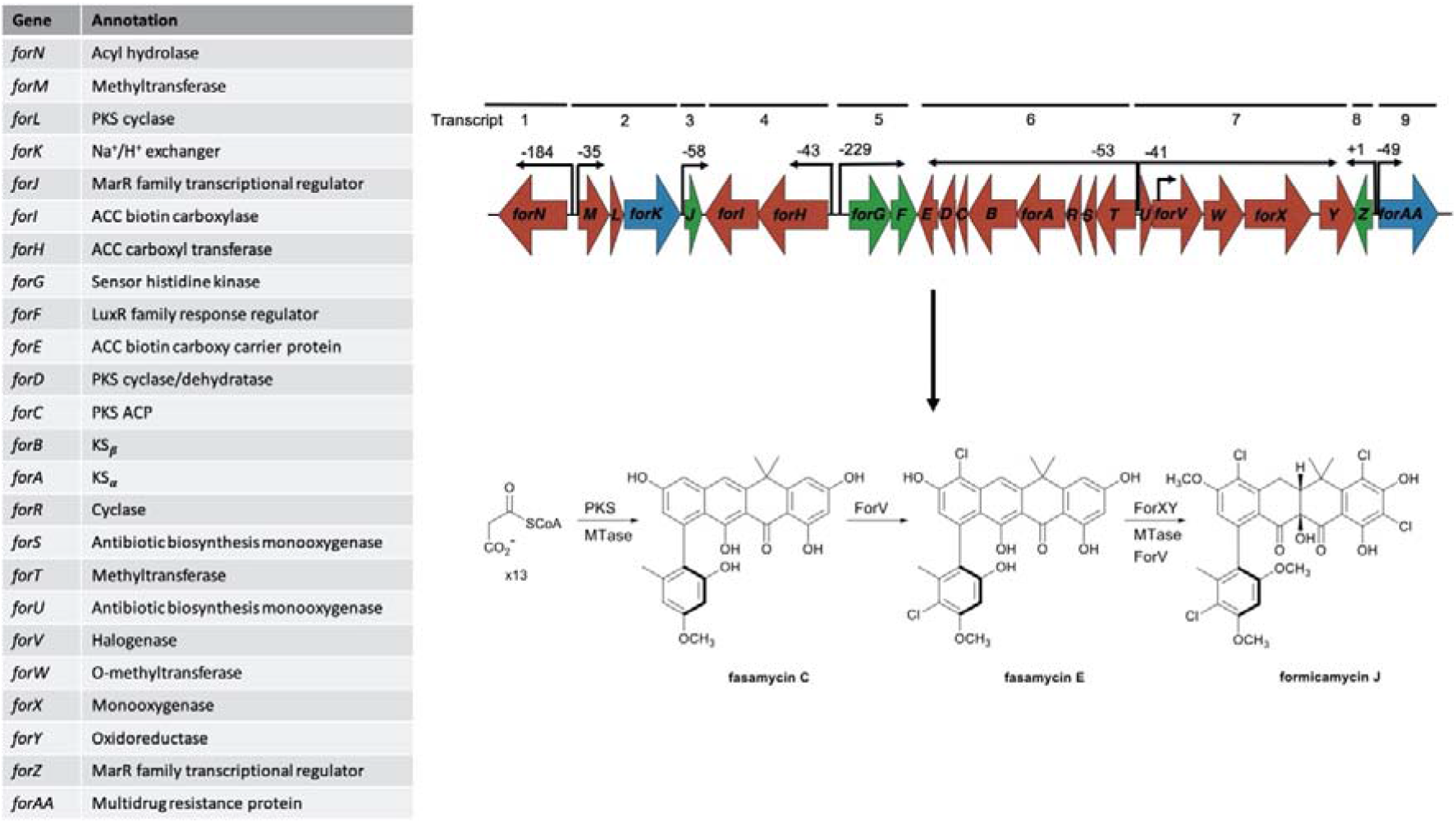
Formicamycin biosynthesis requires 24 genes expressed on nine transcripts. The minimal *for* BGC contains 24 genes required for formicamycin biosynthesis. Cappable RNA-sequencing identified 10 transcription start sites in the *for* BGC, nine of which are in intergenic regions that likely represent promoter regions for the biosynthetic genes. Formicamycin biosynthesis occurs by the formation of fasamycins through the action of the PKS and associated gene products including methyltransferases (MTase) and a halogenase. ForX catalysed hydroxylation and ring-expansion leads to a lactone intermediate which undergoes a reductive ring contraction catalysed by the flavin-dependent oxidoreductase ForY to yield the formicamycin backbone.

## Results

### Formicamycin biosynthesis requires 24 genes expressed on nine transcripts

We previously showed that Cas9-mediated deletion of 46kbp of DNA encompassing the *for* BGC in *S. formicae* abolished formicamycin biosynthesis. Production was restored in *S. formicae Δfor* by introducing pESAC13-215-G PAC which carries the entire *for* BGC plus 40-80kbp of DNA on each side, thus proving this genomic region encodes biosynthesis of these molecules ^7^. ChIP- and dRNA-seq (see below) suggested 24 genes form this cluster and we confirmed this by deleting genes on each side of this 24 gene cluster contained on pESAC13-215-G and demonstrating their ability to induce formicamycin biosynthesis in the *S. formicae Δfor* strain. To elucidate the transcriptional organisation of the *for* BGC we mapped the transcription start sites (TSS) using cappable RNA-sequencing (accession number E-MTAB-7975) and identified 10 TSS, nine of which are in intergenic regions. We thus conclude that the 24 genes required for formicamycin biosynthesis and export are expressed as nine transcripts comprising *forN, forMLK, forJ, forHI, forGF, forTSRABCD, forUVWXY, forZ* and *forAA*. The other TSS is located within the *forV* coding region and likely maintains the expression levels of *forWXY*. The *forX* and *forY* gene products are required for the unique ring conversion that changes a fasamycin precursor into a formicamycin ^13^ (**Figure 1**).

### Formicamycin biosynthesis is repressed by the MarR-family regulator ForJ and activated by the two-component system ForGF

The *for* BGC encodes three CSRs that were predicted to regulate the expression of the transcripts required for formicamycin biosynthesis and export. There are two putative MarR-family transcriptional regulators encoded by *forJ* and *forZ* and a TCS encoded by *forGF*. To determine the roles of these CSRs we made CRISPR/Cas9-mediated deletions in each of their coding genes and measured formicamycin biosynthesis in the resulting mutants. Formicamycin and fasamycin biosynthesis were abolished in the Δ*forGF* mutant, whereas ectopic expression of an additional copy of *forGF* from the native promoter led to an almost doubling (1.9-fold increase) in formicamycin biosynthesis when grown in solid culture. The loss of production caused by deletion of *forGF* was rescued by ectopic expression of *forGF*. In contrast, formicamycin production increased 5-fold in the Δ*forJ* mutant when compared to the wild-type with the levels of fasamycin precursors increasing more than 28-fold. This suggests that the conversion of a fasamycin to a formicamycin is the limiting step in this biosynthetic pathway (the absolute titres of fasamycins are generally lower than those of the formicamycins). Overall, the combined production of fasamycin and formicamycin metabolites increased 6.7-fold in the Δ*forJ* mutant grown in solid culture. Introduction of *forJ* cloned downstream of either the putative *forJ* promoter or the constitutive *ermE\** promoter failed to complement the *ΔforJ* mutant for reasons we cannot explain, but overexpression of *forJ* under control of the *ermE\** promoter in the wild-type strain significantly reduced fasamycin and formicamycin biosynthesis. Deletion of *forZ* reduced formicamycin biosynthesis to approximately 60% of the wild-type levels (Δ70% of total metabolites). Taken together, these results show that formicamycin biosynthesis is activated by ForGF and repressed by ForJ, whilst ForZ may be involved in activation of formicamycin biosynthesis but is not absolutely required and most likely regulates the divergent transporter gene *forAA*. We then deleted both *forJ* and *forGF* in combination and this resulted in a mutant that over-produced formicamycins and accumulated fasamycins, even though loss of *forGF* was expected to result in biosynthesis being abolished. Furthermore, as noted above, the ectopic expression of a second copy of *forJ* under a native promoter is enough to almost abolish biosynthesis in the wild-type strain, even in the presence of the activating TCS. These combined data suggest that *forJ* sits at the top of the *for* BGC regulatory network as the effects of manipulating the other CSRs are overcome by its activity (**Figure 2, Table 1**).

**Table 1:** Fasamycin and formicamycin production by engineered *Streptomyces formicae* strains on solid agar and in liquid culture. Values are mean ± s.d.; Wild-type *n* = 16; mutants *n* = 3; \**forJ* under control of the native promoter; \*\**forJ* under control of the ErmE* promoter.

| Strain | Fasamycin titre (μM) | Formicamycin titre (μM) | Combined titre (μM) | Fasamycins titre (μM) | Formicamycin titre (μM) | Combined titre (μM) |
| --- | --- | --- | --- | --- | --- | --- |
|  | Solid |  |  | Liquid |  |  |
| Wild-type | 6.5 ± 9.6 | 81.3 ± 21.6 | 87.9 ± 74.2 | 0 | 0 | 0 |
| Wild-type + *forJ* | 0 | 24.8 ± 4.7 | 24.8 ± 4.7 | 0 | 0 | 0 |
| Δ*forJ* | 186.2 ± 22.2 | 406.5 ± 42.1 | 592.7 ± 69.4 | 6 ± 0.2 | 624.5 ± 29.4 | 628.4 ± 34.9 |
| $\Delta$ *forJ* + *forJ* * | 170.8 ± 15.4 | 558.8 ± 53.5 | 729.6 ± 74.8 | 2.7 ± 0.6 | 657.3 ± 40.7 | 662.1 ± 47.1 |
| $\Delta$ *forJ* + *forJ* ** | 78.0 ± 60.2 | 784.0 ± 18.9 | 862.0 ± 36.6 | 210 ± 36.0 | 423.1 ± 72.4 | 633.1 ± 138.1 |
| Wild-type + *forGF* | 8.6 ± 0.02 | 155.7 ± 18.7 | 164.3 ± 23.7 | Not tested | Not tested | Not tested |
| Δ*forGF* | 0 | 0 | 0 | Not tested | Not tested | Not tested |
| Δ*forGF* + *forGF* | 13.1 ± 6.7 | 153.7 ± 73.6 | 165.7 ± 86.1 | Not tested | Not tested | Not tested |
| Δ*forJ* + *forGF* | 56.7 ± 52.9 | 814.2 ± 139.8 | 873.4 ± 195.9 | 275.4 ± 11.6 | 759.8 ± 193.8 | 1035.2 ± 212.6 |
| Δ*forJ* Δ*forGF* | 76.3 ± 36.8 | 648.1 ± 112.7 | 724.5 ± 147.8 | 242.9 ± 21.6 | 355.3 ± 82.1 | 598.1 ± 85.7 |
| Δ*forJ* Δ*forZ* | 38.6 ± 31.2 | 388.5 ± 24.8 | 427.1 ± 31.6 | 516.48 ± 297.4 | 409.1 ± 95.4 | 925.6 ± 341.2 |
| Δ*forZ* | 11.1 ± 10.6 | 49.8 ± 3.7 | 60.9 ± 19.7 | Not tested | Not tested | Not tested |
| $\Delta forZ + forZ$ | $13.6 \pm 15.8$ | $21.0 \pm 15.7$ | $31.8 \pm 38$ | Not tested | Not tested | Not tested |
| $\Delta forV$ | $72.8 \pm 60.3$ | 0 | $72.8 \pm 60.3$ | $1.3 \pm 1.8$ | 0 | $1.3 \pm 1.8$ |
| $\Delta forJ\Delta forV$ | $587.5 \pm 268.1$ | 0 | $587.5 \pm 268.1$ | $79.4 \pm 8.1$ | 0 | $79.4 \pm 8.1$ |
| $\Delta forX$ | $42.3 \pm 22.6$ | 0 | $42.3 \pm 22.6$ | $2.54 \pm 4.6$ | 0 | $2.54 \pm 4.6$ |
| $\Delta forJ\Delta forX$ | $782.6 \pm 147.8$ | 0 | $782.6 \pm 147.8$ | $274.3 \pm 325.4$ | 0 | $274.3 \pm 325.4$ |

**Figure 2.**
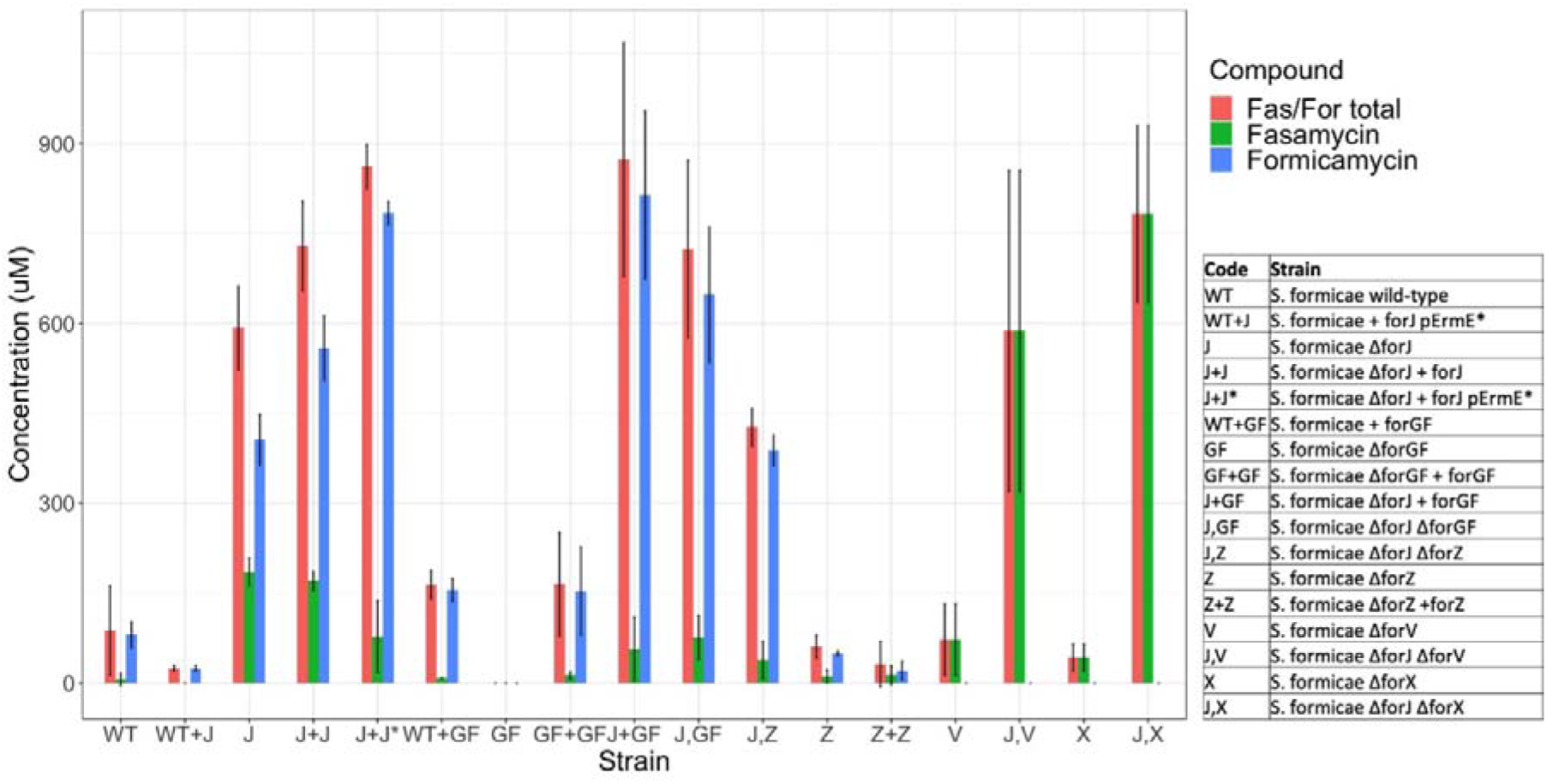
Manipulation of BGC situated regulators affects formicamycin biosynthesis. Deletion of *forJ* results in overproduction of formicamycins and accumulation of the fasamycin precursors. Deletion of *forGF* abolishes fasamycin and formicamycin production. Deletion of *forJ* combined with a second copy of *forGF* results in 10-fold higher formicamycin production than the wild-type strain. Deletion of *forJ* can also be combined with mutations in biosynthetic machinery to generate strains that accumulate precursors and intermediates. Deletion of *forZ* results in a reduction of formicamycin biosynthesis to around 60% of the wild-type strain on solid agar. Manipulation of *forJ* is enough to overcome any other regulatory mutation. Error bars represent standard deviation across experimental replicates. Values are mean ± s.d.; Wild-type *n* = 16; mutants *n* = 3;

To determine how these CSRs control the expression of the *for* biosynthetic genes we generated 3x-Flag-tagged fusion constructs to complement the deletion mutants and used ChIP-sequencing to identify where these CSRs bind across the *S. formicae* chromosome (accession number E-MTAB-8006). The results show that ForF binds to a single site in the *for* BGC at the promoter region between the *forGF* operon and the divergent *forHI* operon, and therefore likely autoregulates and controls expression of *forHI*. ForZ binds to a single site between the divergent *forZ* and *forAA* genes and likely acts as a typical MarR-family regulator by controlling expression of itself and the MFS transporter gene *forAA*. ForJ, the apparent master CSR, binds to multiple locations across the BGC. There are binding sites between the divergent *forN* and *forMLK* operons, in the intergenic regions between the divergent *forHI* and *forGF* operons, and between the *forTSRABCDE* and *forUVWXY* operons that encode the majority of the core biosynthetic machinery. These data are consistent with the observation that ForJ represses the biosynthesis of fasamycins and formicamycins by repressing the transcription of most of the biosynthetic genes. ForJ also binds to its own coding region, presumably to autorepress via a roadblock mechanism, as well as binding within the coding region of *forE* at the end of the long *forTSRABCDE* transcript. We predict that its likely function here is to act as a roadblock to prevent the RNA polymerase transcribing the *forTSRABCDE* operon from running into RNA polymerase transcribing *forGF* since this transcript is required for activation of the *for* BGC (**Figure 3**). Only three significant enrichments occurred outside the *for* BGC: ForF binds upstream of *KY5_0375* which encodes a putative NLP/P60 family protein, while ForJ binds upstream of both *KY5_3182* which encodes a putative MoxR-type ATPase, and *KY5_5812*, which encodes a hypothetical protein. The significance of these binding events is not known, and they were not considered further in this study.

**Figure 3.**
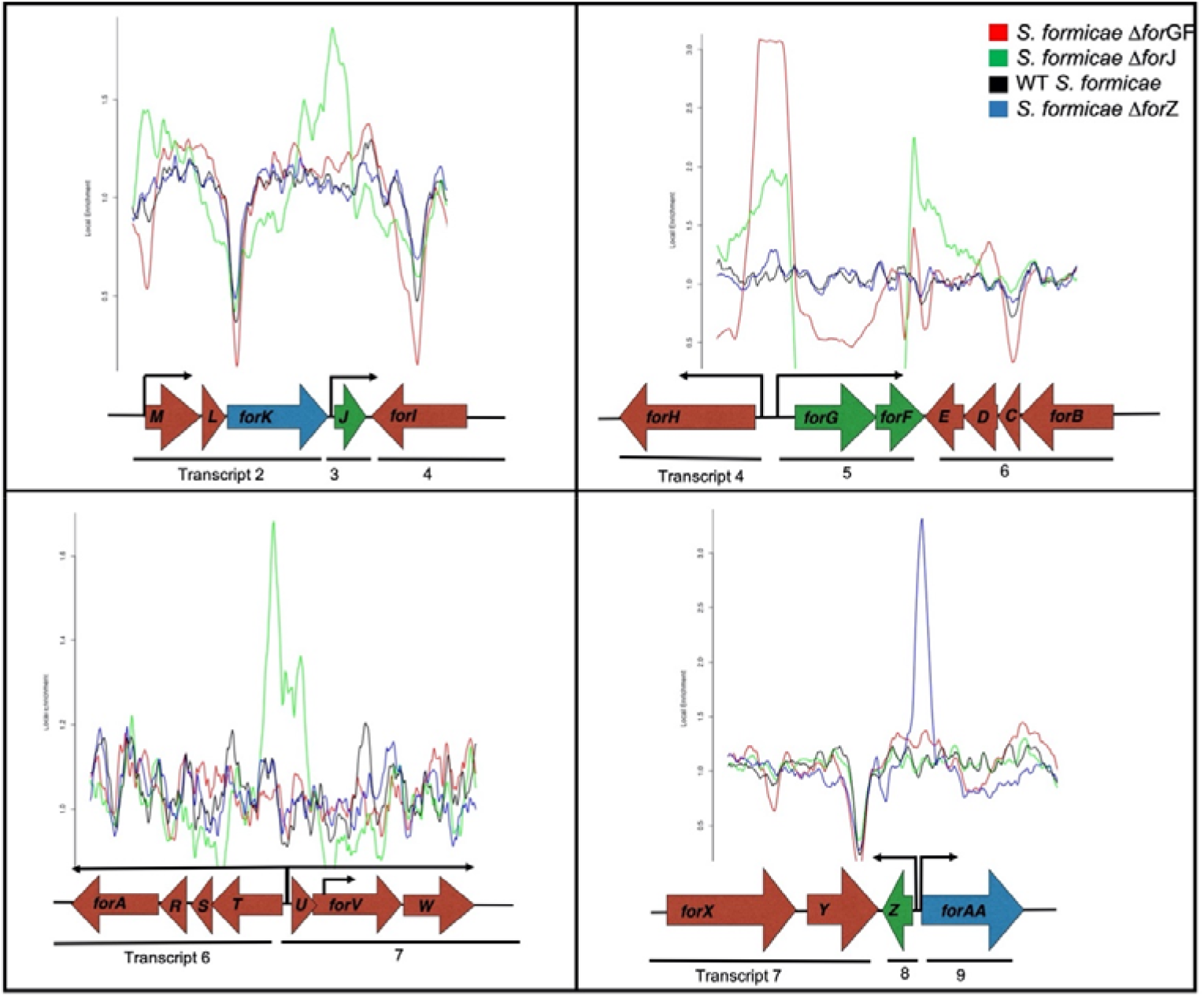
The *for* CSRs bind to multiple promoter regions within the *for* BGC. ForJ binds to multiple locations across the *for* BGC to regulate the expression of the majority of the genes required for formicamycin biosynthesis. ForGF binds to a single promoter within the *for* BGC to regulate itself and the divergent *forHI* transcript. ForZ binds a single promoter between itself and the divergent transporter gene *forAA* and is not predicted to directly regulate biosynthesis of the formicamycins.

To determine the roles of ForGF, ForJ and ForZ in regulating *for* BGC expression we compared the mRNA levels of all *for* transcripts in the Δ*forJ,* Δ*forGF* and Δ*forZ* mutants to those in the wild-type strain, with the exception of *forN* as no suitable primers for qRT-PCR could be found within this relatively short transcript (**Figure 4**). The results show that levels of all transcripts encoding core biosynthetic machinery were higher in the Δ*forJ* mutant compared to the wild-type strain, consistent with the hypothesis that binding of ForJ represses the expression of these transcripts. Since the *forJ* transcript is missing from the Δ*forJ* mutant we made a transcriptional fusion between the *forJ* promoter and *gusA* (encoding β-glucouronidase (GUS)) and found that GUS activity was 2-fold higher in Δ*forJ* relative to the wild-type, suggesting ForJ is autorepressed (*Table S1*). In contrast, levels of the core biosynthetic transcripts were greatly reduced in the Δ*forGF* mutant compared to the wild-type strain which is consistent with the fact that this mutant does not make fasamycins or formicamycins. Activity levels of the *forG* and *forH* promoters was also reduced in the Δ*forGF* mutant, suggesting ForGF auto-activates. Interestingly, levels of the *forJ* transcript were slightly increased in the *forGF* mutant, suggesting levels of repression from *forJ* are higher in the absence of activation by *forGF.* In the Δ*forZ* mutant, levels of the *forAA* transcript were decreased more than three-fold suggesting that ForZ is required to activate the production of this putative transporter. Levels of some of the transcripts encoding biosynthetic machinery were also reduced in the *forZ* mutant compared to the wild-type strain, suggesting that *forZ* may play an indirect role in activating formicamycin biosynthesis (**Figure 4**). Interestingly, the activity of the *forZ* promoter was increased in the Δ*forZ* mutant, suggesting that ForZ may be an example of a dual activator-repressor MarR regulator that activates expression of the divergent transporter while repressing its own transcription. Together these results show that ForJ represses the majority of the core biosynthetic genes by binding to multiple regions across the *for* BGC. We also propose that ForGF is required to activate the divergent *forGF* and *forHI* promoters. The *forHI* genes encode subunits of the acetyl-CoA carboxylase which converts acetyl-CoA into malonyl-CoA, the substrate of the *for* PKS, and are essential for the initiation of formicamycin biosynthesis. We hypothesise that expression of the remaining biosynthetic genes is repressed by ForJ in order to ensure biosynthesis only begins when sufficient levels of malonyl-CoA for formicamycin biosynthesis have been achieved. The results also indicate that ForZ auto-represses while activating the transporter gene *forAA* to ensure that compounds do not accumulate intracellularly once biosynthesis has been initiated.

**Figure 4.**
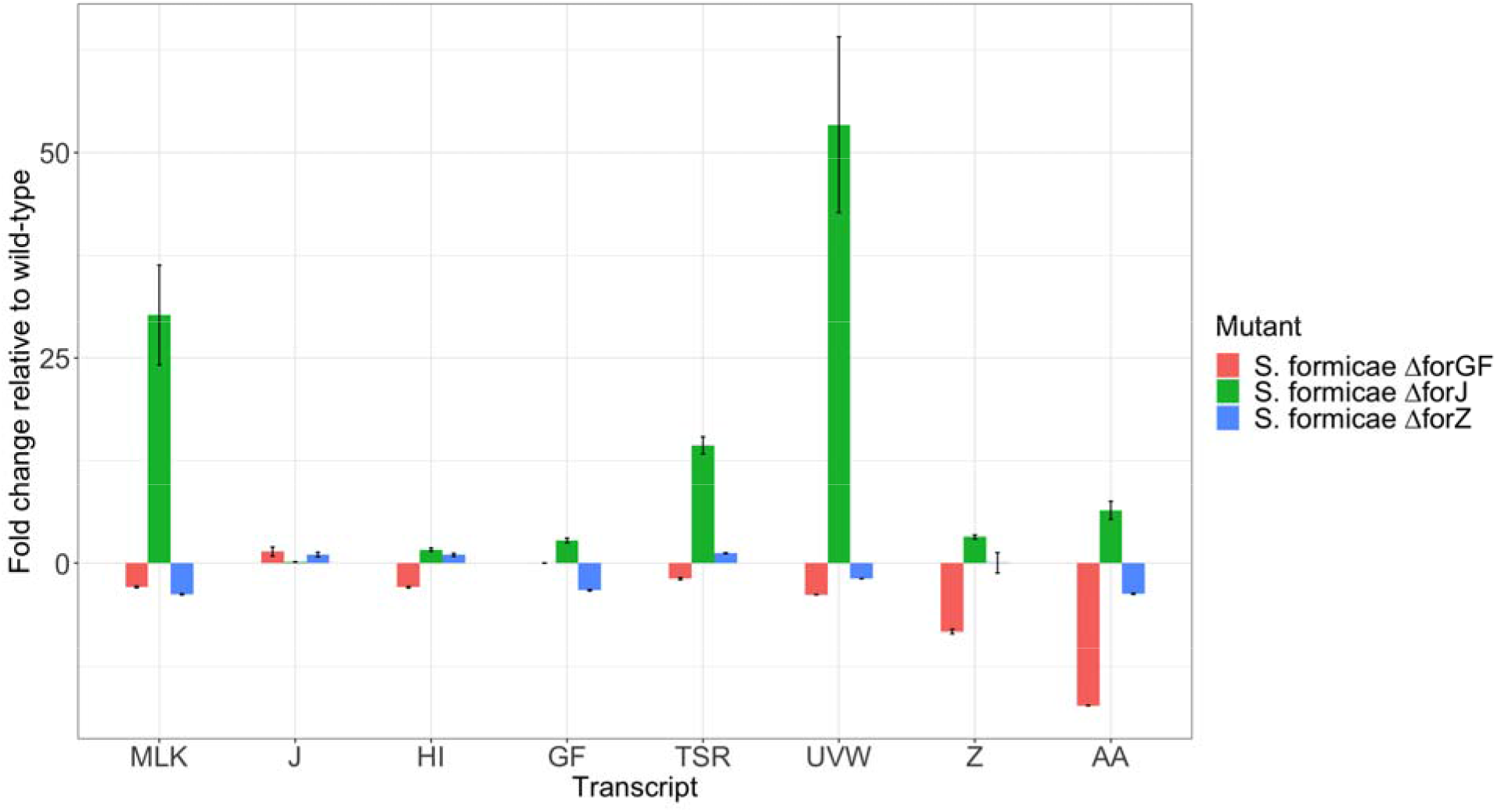
Deletion of *forJ* results in increased transcription of *for* genes and deletion of *forGF* decreases expression. In accordance with observed titres, deletion of *forJ* results in an increase in the expression of all other *for* cluster transcripts. Deletion of *forGF* results in a decrease in expression of all biosynthetic transcripts but an increase in the expression of the repressor gene *forJ*, thereby inhibiting biosynthesis. In the *forZ* deletion mutant, expression of the resistance transporter *forAA* is decreased, suggesting *forZ* is required to activate its transcription. There is a small decrease in the levels of other transcripts in the *forZ* mutant, suggesting this regulator may also indirectly influence expression of these genes without binding to their promoters. Error bars represent standard deviation across experimental replicates.

### BGC derepression results in the production of new formicamycins and induces biosynthesis in liquid culture

Wild-type *S. formicae* does not produce fasamycins or formicamycins during liquid culture and production levels on solid agar are low. This limits the scope for further investigation of the antibiotic potential of these compounds since they cannot easily be purified on a large scale. As noted, the ForGF TCS is essential for biosynthesis in the wild-type while deletion of *forJ* leads to 5-fold higher production of formicamycins (6.7-fold increase in combined fasamycin and formicamycins) on solid medium. We therefore expressed a second copy of the *forGF* operon in the Δ*forJ* strain and found the resulting strain makes 10-fold higher levels formicamycins on solid culture when compared to the wild-type (**Figure 2, Table 1**). Further analysis of the Δ*forJ* and Δ*forJ*+*forFG* strains gave the surprising result that deletion of *forJ* induced production of the formicamycins in liquid medium. The Δ*forJ* mutant produced 1.5-fold more formicamycins when grown in liquid culture compared to on solid culture, although with significantly reduced levels of accumulated fasamycins; this equates to approximately the same overall production level of both sets of metabolite combined, and to an overall 7.1-fold increase in total productivity versus the wild-type strain grown on solid culture. The Δ*forJ*+*forGF* mutant was even more impressive producing 1.6-fold the total levels of metabolites compared to the Δ*forJ* mutant in liquid culture and 11.8-fold more total metabolites (9.3-fold more formicamycins) than the wild-type strain grown in solid culture. By enabling production of the formicamycins during liquid culture these mutants will facilitate fermenter scale production to accelerate their further investigation. Deletion of *forJ* also induced the production of two new formicamycin congeners (formicamycins S and R) each carrying 5 chlorine atoms (**Figure 5**) ^14^. Previously we had observed a maximum of 4 chlorination events for any congener. Both of these molecules exhibited potent antibacterial activity against both methicillin sensitive and resistant strains of *Staphylococcus aureus* (MSSA and MRSA respectively, **Table 2**).

**Table 2:**
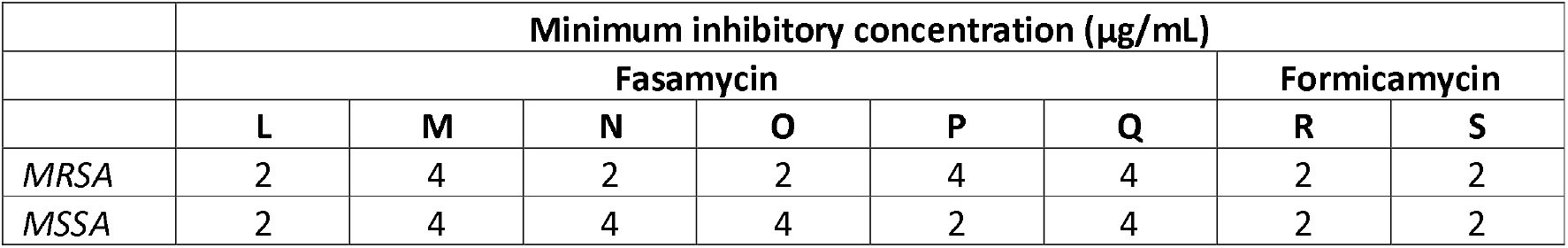
Minimal inhibitory concentration of new fasamycin and formicamycin congeners against *S. aureus* ATCC BAA-1717 (*MRSA*) and ATCC 6538P (MSSA).

|  | Minimum inhibitory concentration (µg/mL) |  |  |  |  |  |  |  |
| --- | --- | --- | --- | --- | --- | --- | --- | --- |
|  | Fasamycin |  |  |  |  |  | Formicamycin |  |
|  | L | M | N | O | P | Q | R | S |
| MRSA | 2 | 4 | 2 | 2 | 4 | 4 | 2 | 2 |
| MSSA | 2 | 4 | 4 | 4 | 2 | 4 | 2 | 2 |

**Figure 5.**
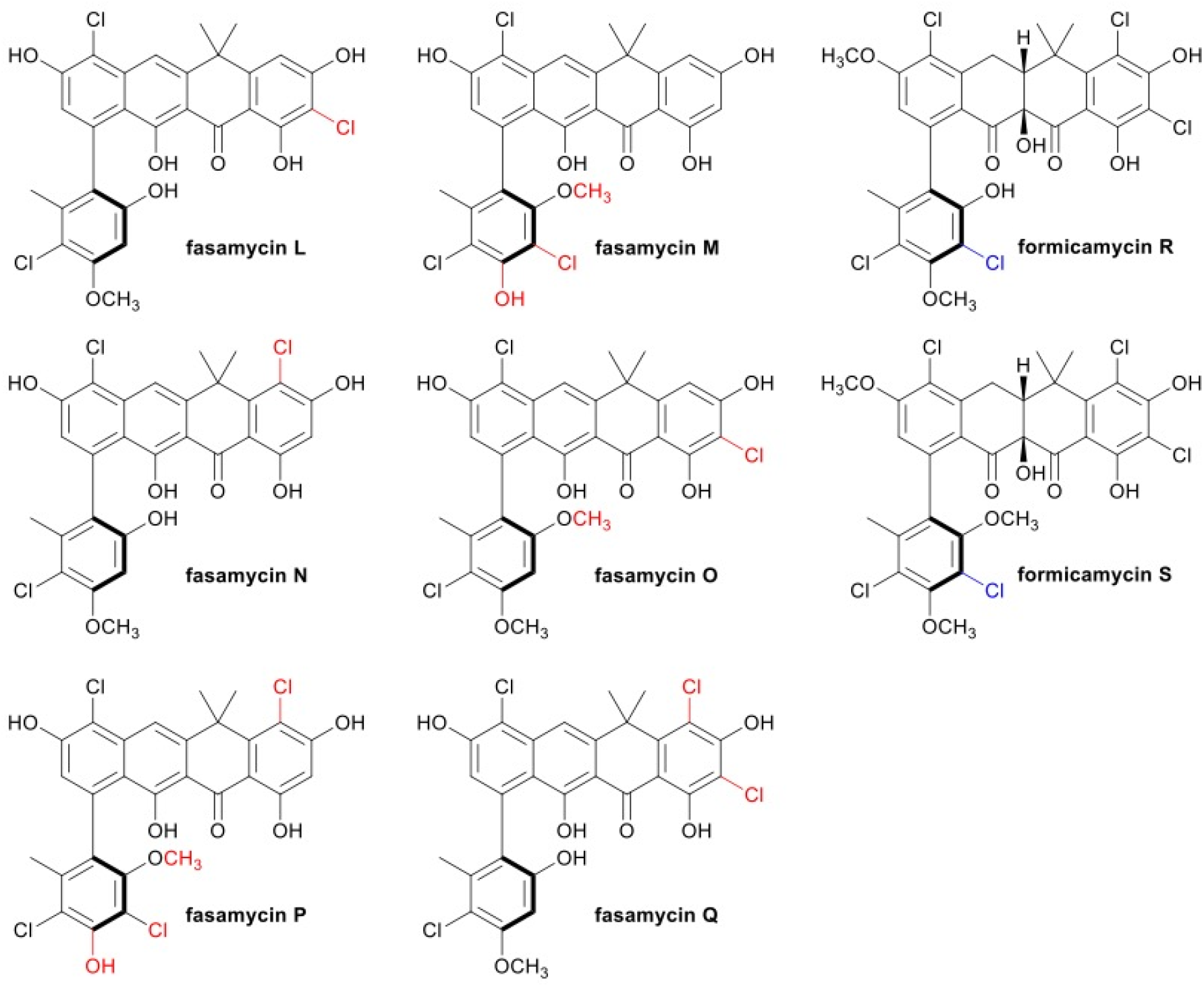
New fasamycin and formicamycin congeners isolated from de-repressed *for* cluster mutants. Deletion of *forJ* results in accumulation of all previously identified congeners from the *for* BGC in addition to as the production of two new formicamycin congeners (R & S) which exhibit a unique C4 chlorination (highlighted in blue). De-repressing the *for* BGC by deleting *forJ* in the *S. formicae ΔforX* mutant results in the production of 6 new fasamycin congeners (L to Q) which have additional chlorination and methylation patterns (highlighted in red) compared to fasamycins C-E produced by the wild-type and *S. formicae ΔforX* strains. All the new congeners displayed potent antibacterial activity against MRSA.

### Combining mutations in regulatory and biosynthetic genes results in the accumulation of new fasamycin congeners

Our observations suggested that combining de-repression (deletion of *forJ*) with deletion in key biosynthetic genes would lead to strains that accumulate elevated levels of pathway intermediates and, potentially, new congeners. We previously showed that the halogenase ForV performs a gatekeeper function, with chlorination of fasamycin intermediates controlling the ability of downstream enzymes to utilise these molecules as substrates and their conversion into the formicamycin skeleton ^7^. We thus constructed a *S. formicae* Δ*forJ*Δ*forV* strain and found that it accumulates fasamycin C exclusively at a titre similar to that of the combined fasamycins and formicamycins produced by the Δ*forJ* strain in solid culture; this corresponds to a titre 90-fold higher than the total fasamycins produced by the wild-type strain (**Table 1, Figure 2**). This provides a powerful route for selectively producing non-halogenated fasamycin congeners, and the lack of formicamycin biosynthesis by this strain is consistent with the proposed gatekeeper function of ForV.

In a similar vein we next made a strain lacking *forJ* and *forX* which encodes the flavin dependant monooxygenase ForX, the enzyme responsible for the first ring-expansion step involved in converting fasamycin precursors into formicamycins ^13^. The resulting Δ*forJ*Δ*forX* strain did not make formicamycins but instead accumulated chlorinated fasamycin congeners to approximately 120-times the level of the wild-type strain (8.9-fold increase in total metabolites) when grown in solid culture (**Table 1, Figure 2**). Moreover, the congener profile of this strain changed considerably and six new fasamycin congeners (**L-Q**) were isolated (**Figure 5**) ^14^. These include molecules carrying up to four chlorine atoms, whereas a maximum of two had previously been observed for fasamycins isolated from the wild-type strain. These new congeners all displayed potent antibacterial activity against both MSSA and MRSA (**Table 2**).

The structures of the new fasamycin and formicamycin congeners were assigned using HRLCMS and 2D NMR based on our published data (Qin et al., 2017) ^7^. The substituent variations in chlorination and O-methylations were determined from the 2D HSQC and NOESY data. Supplementary chemistry data are available here: https://figshare.com/articles/dataset/Structure_determination_of_new_fasamycin_and_formicamycin_congeners/12951974 ^14^.

## Discussion

Using targeted gene deletions and cappable-RNA sequencing we have shown that 24 genes are required for formicamycin biosynthesis and export. These are expressed as nine transcriptional units and are under the control of three CSRs: ForJ and ForZ are MarR-type regulators while ForGF is a TCS. The latter is responsible for activating expression of these transcripts by binding to a single divergent promoter in the *for* BGC and is essential for formicamycin biosynthesis in the wild-type strain but not in a Δ*forJ* deletion mutant. TCSs are one of the major ways in which bacteria sense changes in their environment and the large genomes of *Streptomyces* species generally encode for high numbers of TCSs which enable them to survive in dynamic and sometimes extreme environments ^15^. Due to the availability of genome sequencing data and protocols for genetic manipulation, the majority of characterised TCSs are from model organisms such as *S. coelicolor* and *S. venezuelae* ^16^. Of the TCSs studied in these model species, several have been shown to control secondary metabolism directly, while the majority coordinate secondary metabolism and morphological development either via global regulation (e.g. PhoPR, MtrA/B) or by interacting with CSRs (e.g. AfsQ1) ^17^. In contrast, ForGF is a rare example of a cluster situated TCS that specifically activates transcription of the *for* biosynthetic genes. Another example of a cluster-situated TCS is *cinKR* which, when deleted, abolishes biosynthesis of the lanthipeptide antibiotic cinnamycin much like deletion of *forGF* in *S. formicae* ^18^. However, it should be noted that CinKR is responsible for the activation of a resistance transporter that is absolutely required for cinnamycin biosynthesis, rather than directly activating biosynthetic genes as ForGF appears to. Cluster-situated TCSs that activate biosynthesis in this way represent a promising target for overexpression to activate BGCs that may be cryptic or expressed at low levels like the *for* BGC. Indeed, ectopic expression of an extra copy of *forGF* resulted in a significant increase (2-fold) in the level of formicamycin production compared to the wild-type strain in solid culture.

MarR-family regulators such as ForJ usually repress transcription of their target genes by binding to DNA sequences within promoter regions ^19^. We have shown that ForJ is the major *for* cluster repressor and deletion of *forJ* leads to increased production of multiple pathway products and the induction of biosynthesis during liquid culture of *S. formicae*. Deletion of CSRs is a known method of inducing biosynthesis from cryptic BGCs ^20^, however, to our knowledge, MarR regulators that repress entire biosynthetic pathways in this way are relatively rare. MarR regulators typically bind to intergenic regions to autoregulate their own gene expression and the divergently transcribed gene ^21^. Often, the genes under their control are involved in the control of export of secondary metabolites ^22^. ForZ appears to function in this way: through binding to the *forZ-forAA* intergenic region it autoregulates and controls expression of the divergent *forAA* gene, encoding a putative resistance transporter. Interestingly, ForZ appears to activate transcription of *forAA* rather than repressing it, which is unusual although not unique. ForZ is not required for formicamycin biosynthesis and ChIP-seq data shows that it specifically binds to only a single site within the *for* BGC. However, our data indicate that ForZ may indirectly activate the expression of some *for* BGC transcripts to increase formicamycin biosynthesis.

Based on these data we propose a model in which ForG senses an (unknown) environmental change and phosphorylates ForF, which then activates expression of *forHI* which encode subunits of the acetyl-CoA carboxylase that converts acetyl-CoA to the polyketide precursor malonyl-CoA. MarR-family regulators are also known to bind small molecule ligands and often these ligands are products of the pathways within which they are encoded ^22^. We therefore hypothesise that production of malonyl-CoA above a certain threshold level either directly or indirectly induces ForJ and leads to de-repression of the *for* BGC. This results in the expression of the seven transcripts under the regulation of ForJ which contain the biosynthetic machinery required for formicamycin biosynthesis. To prevent these compounds accumulating intracellularly, we suggest that ForZ activates transcription of forAA to export formicamycins. Where MarR regulators have been shown to regulate an efflux pump, they generally bind and respond to the molecule required for export ^23^. An example is OtrR encoded within the oxytetracycline BGC of Streptomyces rimosus that controls the expression of the divergent transporter gene otrB in response to the presence of oxytetracycline and biosynthetic pathway intermediates^24,25^. It is possible that ForZ is sensitive to fasamycin and/or formicamycin levels and activates expression of forAA to prevent toxic accumulation while ensuring expression of other for transcripts remains low until this resistance mechanism is activated. This will be investigated in future studies.

The formicamycins represent promising candidates for investigation as a new structural class of antibiotics due to their potent bioactivity against drug-resistant pathogens and their high barrier to the development of resistance ^7^. Until now, further investigation into the formicamycins has been hindered by their low production levels in *S. formicae* and the fact they were only produced in solid culture. In this work, we show that de-repression of the *for* BGC not only results in increased titres, but also the induction of biosynthesis during liquid culture. This is significant because the majority of industrial antibiotic production is performed in liquid cultures ^26^. Furthermore, using our knowledge of for regulation and biosynthesis we created a series of targeted mutants by combining mutations in genes encoding the biosynthetic machinery with pathway de-repression. This led to *S. formicae* strains that produce high titres of specific metabolites in both solid and liquid culture. This work will greatly accelerate further investigations into these exciting molecules and demonstrates the importance of studying both the biosynthesis and regulation of BGCs encoding specialised metabolites with antibiotic potential.

## Methods

Chemicals and reagents were laboratory standard grade or above and purchased from Sigma Aldrich (UK) or Thermo Fisher Scientific (UK) unless otherwise stated. All media and solutions were made using deionised water (dH_2_O) except where stated otherwise (**Table 3**).

**Table 3:** Media used in this study

| Media | Recipe (per litre) | Water | pH |
| --- | --- | --- | --- |
| SFM | 20 g soy flour<br>20 g mannitol<br>20 g agar | Tap |  |
| MYM | 4 g maltose<br>4 g yeast extract<br>10 g malt extract<br>+/- 18 g agar | 50:50 Tap:deionised | 7.3 |
| LB | 10 g tryptone<br>5 g yeast extract<br>10 g NaCl (omitted when selecting with Hygromycin)<br>+/- 20 g agar | Deionised | 7.5 |
| 2YT | 16 g tryptone<br>10 g yeast extract<br>5 g NaCl | Deionised | 7.0 |

Generally, *Streptomyces* strains were grown at 30°C and other organisms at 37°C, with shaking at 200 rpm for liquid cultures, unless otherwise stated (**Table S2**). *Streptomyces* spores were harvested from confluent lawns streaked from single colonies using a sterile cotton bud and stored in 1.5 ml 20% glycerol at −80°C. Glycerol stocks were made by resuspending overnight culture in 50:50 LB and glycerol (final concentration 20%). All plasmids and ePACs used in this study are described in **Table S3** and all primers in **Table S4**. Standard DNA sequencing was conducted by Eurofins Genomics using the Mix2Seq kit (Ebersberg, Germany).

### Standard molecular techniques

Genomic DNA and PACs were isolated by resuspending 1 ml overnight culture in 100 μl solution I (50 mM Tris/HCl, pH 8; 10 mM EDTA). Alkaline lysis was performed by adding 200 μl solution II (200 mM NaOH; 1% SDS) followed by 150 μl solution III (3M potassium acetate, pH 5.5). The supernatant was extracted in 400 μl phenol:chloroform:isoamyl alcohol and the upper phase mixed with 600 μl 2-propanol and incubated on ice for precipitation of the DNA. The DNA was pelleted and washed with 200 μl 70% ethanol, air dried and resuspended in sterile dH_2O_. Plasmid DNA was prepared using the QIAprep Spin Miniprep kit (Qiagen) according to the manufacturer’s instructions. All DNA samples were quantified using the Nanodrop 2000 UV-Vis Spectrophotometer and the Qubit assay using the Qubit^®^ fluorimeter 2.0.

PCRs were conducted using either the PCRBIO Taq DNA Polymerase (PCR Biosystems) for diagnostic reactions or the Q5 High-fidelity DNA polymerase for amplification of fragments required for cloning. Both were used according to the manufacturer’s recommendations with a final concentration of 100 nM primers. PCR products were analysed using gel electrophoresis using 1% agarose gels in TBE buffer (90 mM Tris HCl, 90 mM Boric Acid, 2 mM EDTA) with 2 μg/ml ethidium bromide and visualised by UV-light. When required, bands of interest were excised from the gel and the DNA recovered using the QiaQuick Gel Extraction Kit (QIAGEN) according to the manufacturer’s instructions.

For Golden Gate assembly, 100 ng purified backbone was incubated with 0.3 μl insert, 2 μl T4 ligase buffer (NEB), 1 μl T4 ligase (NEB) and 1 μl of the relevant restriction enzyme in a total volume of 20 μl made up in dH_2O_. Reactions were incubated under the following conditions: 10 cycles of 10 minutes at 37°C and 10 minutes at 16°C, followed by 5 minutes at 50°C and 20 minutes at 65°C. Plasmids were digested in 50-100 μl total volumes with restriction enzymes and their appropriate buffer (ether Roche or NEB) in accordance with the manufacturers recommendations (typically 1 μg of DNA was digested with 1 unit of enzyme for 1 hour at 37°C). Restriction enzymes were heat inactivated at 65°C for 10 minutes and 2 μl shrimp alkaline phosphatase was added to single-enzyme digestions to prevent re-ligation. Ligations were carried out using T4 DNA Ligase according to the manufacturer’s recommendations with a standard ratio of 1:3 plasmid:insert. Multiple DNA fragments were assembled into digested vector backbones using Gibson assembly by designing overlaps of between 18 and 24 nucleotides. Gel extracted DNA fragments were incubated in a ratio of 1:3 of plasmid to insert (1:5 for inserts smaller than 300 nucleotides) in the presence of Gibson Assembly master mix (NEB) at 50°C for 1 hour.

*E. coli* was transformed using heat shock (30 seconds at 42°C) for chemically competent cells or electroporation in a BioRad electroporator (200 Ω, 25 μF and 2.5 kV). PACs were moved into *E. coli* using tri-parental mating by incubating 20 μl of each strain in the centre of an LB agar plate at 37°C and then re-streaking the spot onto selective agar for the desired plasmid combination. Plasmids were conjugated into *S. formicae* via the non-methylating *E. coli* ET12567/pUZ8002 as described previously using between 10 and 200 μl of spores depending on the application (more spores for conjugation of pCRISPomyces-2) ^27^.

To analyse protein content, whole cell lysates were incubated at 100°C for 10 minutes in 50 μl SDS loading buffer (950 μl Bio-Rad^®^ Laemmli buffer, 50 μl β-mercaptoethanol) and analysed by gel electrophoresis on a standard resolving gel of 16% (w/v) acrylamide:Bis-Acrylamide 37.5:1 (Fisher BioReagents). To confirm expression of tagged proteins, proteins were analysed by immunoblot by transferring to a nitrocellulose membrane (Pall Corporation) in a Trans-Blot transfer cell (10 V, 1 hour). After blocking in 5% (w/v) fat-free skimmed milk powder in TBST (50 mM Tris Cl pH 7.5, 150 mM NaCl, 1% Tween) overnight at room temperature, 20 ml HRP-conjugated anti-FLAG antibody diluted 1 in 20000 in TBST was added to the membrane for 1 hour. The membrane was washed 3 times for 10 minutes in TBST before developing in a 50:50 mix of 100 mM Tris pH 8.5, 100 μl luminal, 45 μl coumaric acid and 100 mM Tris pH 8.5, 6 μl 30% hydrogen peroxide for imaging using the ECL setting on a SYNGENE G:Box.

### Editing pESAC-13 to define the borders of the *for* BGC

Targeted mutagenesis of pESAC-13 was conducted by ReDirect as described previously ^27^ except that the *oriT* was removed from the apramycin cassette to avoid undesired recombination events with the PAC. The truncated cassette was PCR amplified, gel purified and electroporated into *E. coli* BW25113 pIJ1790 transformed (by tri-parental mating) with pESAC-13 215-G (containing the *for* BGC). The PCR-confirmed edited PAC was isolated and conjugated into *S. formicae* Δ*for* and the resulting mutants were grown under formicamycin producing conditions (as above). The metabolites were extracted as described below and then analysed by chromatography over a Phenomenex Gemini reversed-phase column (C18, 100 Å, 150×2.1 mm) using an Agilent 1100 series HPLC system and eluting with the following gradient method: 0–2 min 50% B; 2–16 min 100% B; 16–18 min 100% B; 18–18.1 min 100– 50% B; 18.1–20 min 50% B; flowrate 1 mL min^−1^; injection volume 10 μL; mobile phase A: water+0.1% formic acid; mobile phase B: methanol. UV absorbance was monitored at 250, 285, 360 and 415 nm.

### Cappable RNA sequencing

Samples for RNA-sequencing were crushed in liquid nitrogen using a sterile pestle and mortar on dry ice and resuspended by vortexing in 1 mL RLT Buffer (Qiagen) supplemented with β-mercaptoethanol (10 μl in every 1 ml buffer). Following lysis in a QIA-shredder column (Qiagen) the sample was extracted in 700 μl acidic phenol-chloroform and the upper phase mixed with 0.5 volumes 96% ethanol. RNA was purified using the RNeasy Mini spin column (Qiagen) according to the manufacturers protocol, including on column DNase treatment. Following elution, the Turbo-DNase kit was used according to the manufacturers protocol and a further Qiagen RNeasey mini clean-up was conducted before samples were flash frozen in liquid nitrogen for storage at −80°C. Once quantified by Nanodrop and gel electrophoresis, RNA was sent to Vertis Biotechnologie (Freising, Germany) for sequencing and analysed by capillary electrophoresis to map transcription start sites.

### Generating *S. formicae* mutants

Gene deletions were made using the pCRISPomyces-2 system as described previously ^7,12,28^. Approximately 20 bp protospacers were designed so that the last 15 nucleotides, including the PAM (NGG) were unique in the genome. These were annealed and assembled into the sgRNA using a Golden Gate reaction with BbsI and pCRISPomyces-2. The resulting plasmid was then digested with XbaI to allow for the insertion of the repair template (usually approximately 1 Kb from either side of the deletion) using Gibson assembly. Once confirmed by PCR and sequencing, the final vector was conjugated into *S. formicae* via the non-methylating *E. coli* ET12567/pUZ8002. The deletion was confirmed using PCR to amplify the region across the repair template and into the surrounding genomic DNA. Once confirmed, loss of the editing plasmid was encouraged by repeated re-streaking on plates lacking the antibiotic selection and incubating at 37°C.

Complementation of gene deletions were achieved by fusing the gene to either a native promoter in pMS82^29^ by Gibson assembly or by ligating the digested gene product downstream of the constitutive *ermE\** promoter in pIJ10257 ^30^ and transferring the plasmid into the relevant mutant by conjugation.

### ChIP-sequencing

For sampling, spores were inoculated onto cellophane discs and grown for 2, 3 or 4 days. These time points were chosen as day 5 is the earliest formicamycins have been observed in the culture extract and expression of regulator genes would be predicted to happen before biosynthesis. Day 2 was the earliest point that enough biomass could be harvested. Expression of the genes of interest was also confirmed using RT-PCR using the OneStep RT-PCR Kit (Qiagen) before the experiment was conducted. At sampling, discs were removed and the mycelium submerged in 10 ml 1% (v/v) formaldehyde for 20 minutes, followed by 10 ml 0.5 M glycine for 5 minutes. After washing the discs in 25 ml ice-cold PBS (pH 7.4), the samples were frozen at −80°C for storage and a small aliquot retained for immunoblot analysis to confirm expression of the tagged proteins in each sample.

For chromatin immunoprecipitation, cell pellets were resuspended in 2 ml lysis buffer (10 mM Tris-HCl pH 8.0, 50 mM NaCl, 10 mg/ml lysozyme, EDTA-free protease inhibitor) and incubated at 37°C for 30 minutes. To fragment the DNA, 1 ml IP buffer (100 mM Tris-HCl pH 8.0, 500 mM NaCl, 1% v/v Triton-X, EDTA-free protease inhibitor) was added and samples sonicated 20 times at 50 Hz for 10 seconds per cycle, ensuring cool-down on ice for at least 2 minutes between pulses. A small sample of this crude lysate was extracted with phenol:chloroform, treated with RNaseA and analysed by gel electrophoresis to confirm DNA fragments were within the desired size range for sequencing. The remaining crude lydate was cleared by centrifugation and incubated with 40 μl prepared Anti-FLAG M2 beads (Sigma-Aldridge) with rotation at 4°C overnight. To elute the bound DNA, the samples were then incubated in 100 μl elution buffer (50 mM Tris-HCl pH 8.0, 10 mM EDTA, 15 SDS) overnight at 65°C. Another 50 μl elution buffer was then added and the samples incubated for a further 5 minutes at 65°C before DNA from the total 150 μl eluate was purified by adding proteinase K and incubating at 55°C for 1.5 hours. DNA was extracted in 150 μl phenol-chloroform and purified on a QIAquick column (Qiagen) and eluted in 50 μl EB buffer (10 mM Tris-HCl pH 8.5) for quantification and sequencing by Genewiz (Leipzig, Germany) using the Illumina HiSeq platform. Data analysis was conducted as described previously ^31–33^.

### qRT-PCR

RNA samples were confirmed to be DNA-free by conducting test PCRs on the 16S rRNA gene and converted to cDNA using the LunaScript RT SuperMix Kit (NEB) according to the manufacturer’s instructions. Primers for each transcript were optimised using serial dilutions of ePAC template DNA and checked for specificity by melt-curve analysis and gel electrophoresis. Reactions were run in biological triplicate and technical duplicate using the Luna Universal qPCR Master Mix according to the manufacturer’s guidelines in a total volume of 20 μl with approximately 100 ng template cDNA and a final concentration of 0.25 μM of each primer. ΔCT values were normalised to the 16S rRNA gene.

### GUS assay

Strains were generated using pMF96 plasmids containing the *gusA* gene under the control of each relevant promoter, empty pMF96 as a negative control and the pMF23 (*ermEp\**-*gusA*) plasmid as a positive control ^34^ (**Tables S1** and **S3**). GUS plasmids were conjugated into to *S. formicae* WT, *ΔforJ*, *ΔforGF* and *ΔforZ* strains via ET12567/pUZ8002. Three biological replicates of each GUS-producing strain and controls were then confirmed by PCR.

GUS strains were grown on SFM agar with a cellophane disk for 4 days then mycelium was harvested by scraping with a sterile metal spatula and resuspended in 1 mL dilution buffer (50 mM phosphate buffer, 0.1 % Triton X-100 [v/v], 5mM DTT). Samples were sonicated in 3 rounds of 8-10 seconds at 80 Hz before centrifugation at 13,000 rpm, 4 °C for 10 minutes. A total of 95 μL of each sample supernatant was added to a 96 well plate and freeze-thawed at −80 °C and 37 °C, respectively. As strains were grown on agar plates rather than in liquid culture, the protein concentration of lysates were quantified by measuring absorbance at 280 nm rather than measuring cell density at 600 nm. The hydrolysis of PNPG by β-glucuronidase produces galactose and chromophoric 4-nitrophenol (PNP) with a peak absorbance at 420 nm. The intensity of this peak is dependent upon the quantity of enzyme present which correlates with transcription at the inserted promoter. PNPG was added to each well (final concentration 3 μg/mL) to initiate the reaction. Samples were loaded into a plate reader at 37 °C for optimum enzyme activity and quantified at 415 nm (due to some interference with fasamycin peak absorbance at 419 nm) and 550 nm every 5 minutes for 40 minutes, inclusive. Softmax^®^ Pro7 used to extract raw data which was used calculate Miller units/ mg protein (1 unit = 1000*[Abs415–{1.75*Abs550}]/{t*v*Abs280}]). Each promoter’s transcriptional activity was tested in both biological and technical triplicate and mean Miller Units/mg protein calculated. All statistical analysis was performed using IBM SPSSTM Statistics 23.

### Fasamycin and formicamycin congener analysis

#### Solid culture

*S. formicae* WT (*n* = 16) and mutant strains (*n = 3*) were grown on soya flour mannitol (SFM) agar at 30°C for 10 days. Equal size agar plugs (1 cm^3^) were taken in triplicate from each plate and shaken with ethyl acetate (1 mL) for 1 hour before being centrifuged at 3100 rpm for 5 minutes. Ethyl acetate (300 μl) was transferred to a clean tube and solvent was removed under reduced pressure. The resulting extract was dissolved in methanol (200 μL) before being analysed by HPLC (Agilent 1290 UHPLC). To verify peak identity a representative set of samples were analysed by LCMS (Shimadzu IT-ToF LCMS platform). Chromatography was undertaken for both HPLC and LCMS analysis using the following method: Phenomenex Gemini NX C18 column (150 × 4.6 mm); mobile phase A: water + 0.1% formic acid; mobile phase B: methanol. Elution gradient: 0–2 min, 50% B; 2–16 min, 50–100% B; 16–18 min, 100% B; 18–18.1 min, 100–50% B; 18.1–20 min, 50% B; flow rate 1 mL min^−1^; injection volume 10 μL.

#### Liquid culture

*S. formicae* WT or mutant strains (*n* = 3) were grown in liquid TSB (10 mL) for 2 days. 100 μL of 2-day liquid culture was aliquoted into 10 mL of SFM liquid media into sterile 50 mL falcon tubes with sterile bungs. Strains were incubated at 30°C with shaking at 250 rpm. After 10 days, an aliquot (1 mL) of culture was removed from each sample in triplicate and shaken with ethyl acetate (1 mL) for 1 minute and then centrifuged at 3100 rpm for 5 minutes. Ethyl acetate extract (300 μl) was transferred to a clean tube and solvent was removed under reduced pressure. The resulting extract was dissolved in methanol (200 μl) before being analysed by HPLC and LCMS as described before.

#### Titre determination

Titres of fasamycin and formicamycins were determined by comparing peak areas from the above HPLC analysis to those of standard calibration curves and correcting for the concentration that occurred during the extraction process. Calibration curves were determined using standard solutions of fasamycin E (10, 20, 50, 80 and 200 μM) and formicamycin I (10, 20, 50, 100, 200 and 400 μM) (**Figure 6**). The content of fasamycin E and formicamycin I were determined by UV absorption at 418 nm and 285 nm respectively. Each standard solution was measured three times.

**Figure 6.**
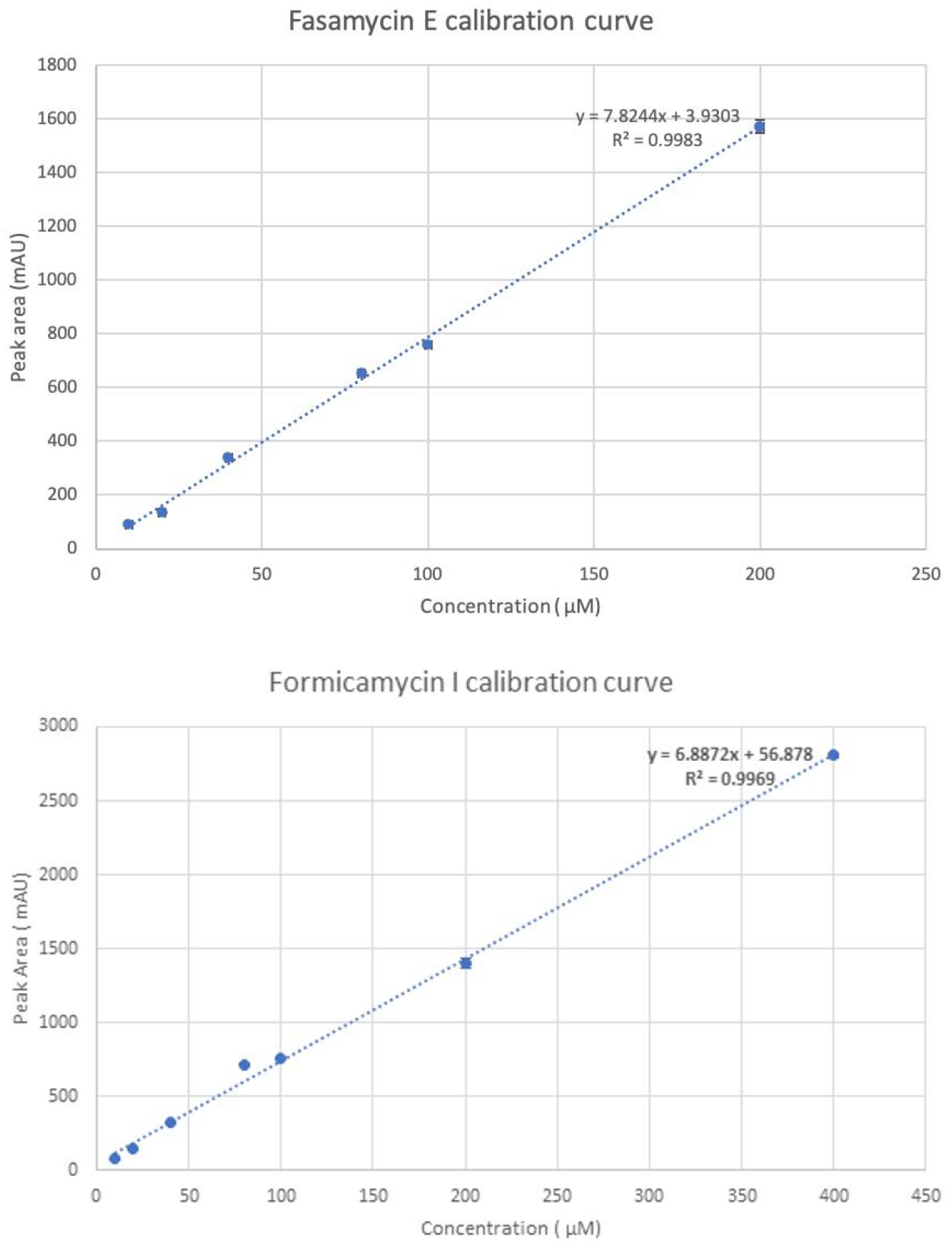
Calibration curve for Fasamycin E and Formicamycin I

### Scale up fermentation of *S. formicae* Δ*forJ* and Δ*forJX*

A seed culture was prepared by inoculating the fresh mycelium from an MS agar plate into TSB liquid medium (50 mL) and incubating for 24 h at 30 °C with shaking at 250 rpm. The seed culture was then used to inoculate TSB (3 L; 6 × 2.5 L flasks, 1:500 inoculation). The cultures were incubated for seven days under the same condition as above after which the whole culture broth was extracted twice with equal amount of ethyl acetate. The organic phase was separated, and the solvent removed by evaporation under reduced pressure. The resulting material was dissolved in methanol and first analysed by LCMS and then subjected to semi-prep purification. In this way six new fasamycin congeners (fasamycins L to Q) were isolated from the Δ*forJX* mutant, and two new formicamycin congeners (formicamycins R and S) from the Δ*forJ* mutant. The structures of these compounds were determined using HRLCMS and 2D NMR (data in ref 14).

#### Analytical LCMS method

For LCMS following analytical LCMS method was used: Phenomenex Kinetex C18 column (100 ÅΔ 2.1 mm, 100 Å); mobile phase A: water +0.1% formic acid; mobile phase B: acetonitrile. Elution gradient: 0–1 min, 20% B; 1–8 min, 20– 100% B; 8–11 min, 100% B; 11–11.1 min, 100–20% B; 11.1–12 min, 20% B; flow rate 0.6 mL min^−1^; injection volume 10 μL.

#### Semi-prep HPLC method for isolation of formicamycins R & S

Chromatography was achieved over a Phenomenex Gemini-NX semi-prep reversed-phase column (C18, 110 Å, 150 ÅΔ 10 mm) using an Agilent 1100 series HPLC system and eluting with the following gradient method: 0–2 min 60% B; 2–18 min 60–100% B; 18–23 min 100% B; 23–23.1 min 100–60% B; 23.1–26 min 60% B; flowrate 3.5 mL min^−1^; injection volume 20 μL; mobile phase A: water +0.1% formic acid; mobile phase B: acetonitrile. The UV absorbance was monitored at 250 and 285 nm.

#### Semi-prep HPLC method for isolation of fasamycins L to Q

Chromatography was achieved over a Phenomenex Gemini-NX semi-prep reversed-phase column (C18, 110 Å, 150 ÅΔ 10 mm) using an Agilent 1100 series HPLC system and eluting with the following gradient method: 0–2 min 70% B; 2–18 min 70–100% B; 18–20 min 100% B; 20–20.1 min 100–70% B; 20.1–24 min 70% B; flowrate 3.5 mL min^−1^; injection volume 100 μL; mobile phase A: water + 0.1% formic acid; mobile phase B: acetonitrile. UV absorbance was monitored at 250, 285, 360 & 415 nm.

### Bioassays

Resazurin assays were performed to determine minimum inhibitory concentrations. A stock solution of compound was prepared in DMSO and diluted in LB or TSB media to give a concentration range of 256 μg/ml to 1 μg/ml containing 5% DMSO. Positive control (PC) for preparations in LB and TSB was apramycin at 50 μg/ml. Negative control (NC) and media control (MC) contained media (LB or TSB) and DMSO at 5%. Methods were followed as detailed in Heine *et al* ^35^. In brief, cultures were grown to confluence overnight and then diluted 1/100 in fresh media and grown to 0.4 OD600nm, diluted to match a 0.5 McFarlands standard and further diluted 1/100. 5 μl of culture was then aliquoted into each well of the 96 well plate excluding the MC well. Plates were grown at 37 °C before being inoculated with 5 μl resazurin dye (6.75 mg / ml, Sigma Aldrich). Colorimetric results were observed 4 hours after inoculation. MSSA (ATCC 6538P) and MRSA (ATCC BAA-1717) indicator strains were obtained from the American Type Culture Collection.

## Acknowledgements

This work was supported by the Biotechnology and Biological Sciences Research Council (BBSRC) via BBSRC Responsive Mode Grants (BB/S00811X/1 and BB/S009000/1) to M.I.H and B.W. (R.D and C.A), and via Institute Strategic Program Project BBS/E/J/000PR9790 to the John Innes Centre (ZQ). HM and KN were supported by BBSRC PhD studentships (BBSRC Doctoral Training Programme grant BB/M011216/1). We thank our colleague Dr Tung Le (John Innes Centre) for useful discussions.

## Author Contributions

R.D, H.M, Z.Q, C.A, K.N, B.W and M.I.H designed the research. R.D, H.M, K.N, B.W and M.I.H wrote the paper and all authors commented. R.D, H.M, K.N and C.A performed the molecular genetics experiments, H.M and Z.Q performed the chemical experiments. G.C conducted the bioinformatics analysis.

