## Supplementary information for "Refactoring the formicamycin biosynthetic gene cluster to make high-level producing strains and new molecules"

**Supporting information**

**Chemistry data is available on Figshare:** <https://figshare.com/articles/dataset/Structure_determination_of_new_fasamycin_and_formicamycin_congeners/12951974>

**GUS assay results from *for* BGC promoters**

**Table S1:** The activities of *for* gene cluster promoters were measured by fusing the promoters upstream of the β-glucoronidase reporter gene in pMF96. The negative control is the empty pMF96 vector with no promoter driving GUS, the positive control plasmid pMF23 has GUS under the control of the high level *ermE** promoter. β-glucoronidase activity was measured by hydrolysis of PNPG using absorbance at 420 nm.

| **Strain Name** | **Mean Miller Units/ mg protein ± Standard Error** |
| --- | --- |
| *S. formicae* WT pMF96 | 0.37 ± 1.10 |
| *S. formicae* WT pMF23 | 91.75 ± 21.75 |
| *S. formicae* *ΔforJ* pMF96 | 0.00 ± 1.67 |
| *S. formicae* *ΔforJ* pMF23 | 86.98 ± 18.81 |
| *S. formicae* *ΔforGF* pMF96 | 2.01 ± 2.49 |
| *S. formicae* *ΔforGF* pMF23 | 100.83 ± 12.22 |
| *S. formicae* *ΔforZ* pMF96 | 0.37 ± 0.56 |
| *S. formicae ΔforZ* pMF23 | 94.44 ± 16.37 |
| *S. formicae* WT p*forJ* | 32.82 ± 12.83 |
| *S. formicae ΔforJ* p*forJ* | 84.85 ± 10.05 |
| *S. formicae* WT p*forG* | 50.56 ± 14.34 |
| *S. formicae* *ΔforGF* p*forG* | 26.68 ± 8.11 |
| *S. formicae* WT p*forH* | 33.62 ± 12.52 |
| *S. formicae* *ΔforGF* p*forH* | 11.86 ± 5.07 |
| *S. formicae* WT p*forZ* | 0.38 ± 1.65 |
| *S. formicae ΔforZ* p*forZ* | 11.41 ± 7.68 |

**Table S2:** Bacterial strains used and generated in this study

| **Strain** | **Description/Genotype** | **Plasmid** | **Resistance** | **Reference/Source** |
| --- | --- | --- | --- | --- |
| *E. coli* Top10 | F– mcrA Δ(mrr-hsdRMS-mcrBC) Φ80lacZΔM15 ΔlacX74 recA1 araD139 Δ(ara leu) 7697 galU galK rpsL (StrR) endA1 nupG |  |  | Invitrogen^TM^ |
| *E. coli* BW25113 | λ^-^, Δ(*araD-araB*)*567,* Δ*lacZ4787*(*::rrnB-4*)*, lacIp-4000*(*lacIQ*), *rpoS369*(*Am*)*, rph-1,* Δ(*rhaD-rhaB*)*568, hsdR514* | pIJ790 | Cml^R^ | ^1^ |
| *E. coli* ET12567 | *dam^-^ dcm^-^ hsdS^-^* | pUZ8002 | Cml^R^/Tet^R^ | ^2^ |
| *E. coli* BL21 | *fhuA2 [lon] ompT gal (λ DE3) [dcm] ∆hsdSλ DE3 = λ sBamHIo ∆EcoRI-B int::(lacI::PlacUV5::T7 gene1) i21 ∆nin5* |  |  | ^3^ |
| *S. formicae* wild-type |  |  |  | Lab stock |
| *MSSA* | ATCC 6538P |  |  | American Type Culture Collection |
| *MRSA* | ATCC BAA-1717 |  |  | American Type Culture Collection |
| *S. formicae ΔforJ* | *forJ* deletion strain |  |  | This work |
| *S. formicae ΔforGF* | *forGF* deletion strain |  |  | This work |
| *S. formicae ΔforZ* | *forZ* deletion strain |  |  | This work |
| *S. formicae ΔforJ: ΦBT1 forJ pforM* | *forJ* complementation under forM promoter | pRD030 | Hyg^R^ | This work |
| *S. formicae ΔforJ: ΦBT1 forJ* | *forJ* complementation under forJ promoter | pRD063 | Hyg^R^ | This work |
| *S. formicae ΔforJ: ΦBT1 pErmE* forJ* | *forJ* complementation under ErmE* promoter | pRD06 | Hyg^R^ | This work |
| *S. formicae ΔforGF: ΦBT1 forGF* | *forGF* complementation under native promoter | pRD031 | Hyg^R^ | This work |
| *S. formicae ∆for:* ΦC31 *for* *1-4* *aac(3)IV* | Whole *for* cluster deletion complemented with pESAC-13 215-G with genes 1-4 (as annotated by AntiSMASH) replaced with apramycin gene | pRD037 | Apr^R^ | This work |
| *S. formicae ∆for:* ΦC31 *for* *1-7* *aac(3)IV* | Whole *for* cluster deletion complemented with pESAC-13 215-G with genes 1-7 replaced with apramycin gene | pRD038 | Apr^R^ | This work |
| *S. formicae ∆for:* ΦC31 *for 32-43* *aac(3)IV* | Whole *for* cluster deletion complemented with pESAC-13 215-G with genes 32-43 replaced with apramycin gene | pRD039 | Apr^R^ | This work |
| *S. formicae ∆for:* ΦC31 *for 36-43 aac(3)IV* | Whole *for* cluster deletion complemented with pESAC-13 215-G with genes 36-43 replaced with apramycin gene | pRD040 | Apr^R^ | This work |
| *S. formicae ∆forJ: ΦBT1 forJ 3x Flag* | *forJ* deletion mutant complemented in-trans with 3x flag-tagged *forJ* for ChIP | pRD034 | Hyg^R^ | This work |
| *S. formicae ∆forGF: ΦBT1 forGF 3x Flag* | *forGF* deletion mutant complemented in-trans with 3x flag-tagged *forGF* for ChIP | pRD035 | Hyg^R^ | This work |
| *S. formicae ∆forZ: ΦBT1 forZ 3x Flag* | *forZ* deletion mutant complemented in-trans with 3x flag-tagged *forZ* for ChIP | pRD036 | Hyg^R^ | This work |
| *S. formicae ΔforV* | *forV* deletion strain |  |  | ^4^ |
| *S. formicae ΔforX* | *forX* deletion strain |  |  | ^5^ |
| *S. formicae ΔforY* | *forY* deletion strain |  |  | ^5^ |
| *S. formicae ΔforS* | *forS* deletion strain |  |  | ^6^ |
| *S. formicae ΔforJ, ΔforV* | *forV* deletion strain with *forJ* deletion |  |  | This work |
| *S. formicae ΔforJ, ΔforX* | *forX* deletion strain with *forJ* deletion |  |  | This work |
| *S. formicae ΔforJ, ΔforY* | *forY* deletion strain with *forJ* deletion |  |  | This work |
| *S. formicae ΔforJ, ΔforS* | *forS* deletion strain with *forJ* deletion |  |  | This work |
| *S. formicae: pMF96* | Wildtype strain with GUS but no promoter controlling expression (negative control) | pMF96 | Hyg^R^ | This work |
| *S. formicae: pMF23* | Wildtype strain with GUS but no promoter controlling expression (negative control) | pMF23 | Apr^R^ | This work |
| *S. formicae ΔforJ: pMF96* | *forJ* deletion strain with GUS but no promoter controlling expression (negative control) | pMF96 | Hyg^R^ | This work |
| *S. formicae ΔforJ: pMF23* | *forJ* deletion e strain with GUS but no promoter controlling expression (negative control) | pMF23 | Apr^R^ | This work |
| *S. formicae ΔforGF: pMF96* | *forGF* deletion strain with GUS but no promoter controlling expression (negative control) | pMF96 | Hyg^R^ | This work |
| *S. formicae ΔforGF: pMF23* | *forGF* deletion e strain with GUS but no promoter controlling expression (negative control) | pMF23 | Apr^R^ | This work |
| *S. formicae ΔforZ: pMF96* | *forZ* deletion strain with GUS but no promoter controlling expression (negative control) | pMF96 | Hyg^R^ | This work |
| *S. formicae ΔforZ: pMF23* | *forZ* deletion e strain with GUS but no promoter controlling expression (negative control) | pMF23 | Apr^R^ | This work |
| *S. formicae ΔforJ: ΦBT1 GUS pforJ* | *forJ* deletion strain with GUS expressed under *pforJ* | pRD062 | Hyg^R^ | This work |
| *S. formicae: ΦBT1 GUS pforJ* | Wildtype strain with GUS expressed under *pforJ* | pRD062 | Hyg^R^ | This work |
| *S. formicae: ΦBT1 GUS pforG* | WIldtype strain with GUS expressed under *pforG* | pRD054 | Hyg^R^ | This work |
| *S. formicae ΔforGF: ΦBT1 GUS pforG* | *forGF* deletion strain with GUS expressed under *pforG* | pRD054 | Hyg^R^ | This work |
| *S. formicae: ΦBT1 GUS pforH* | Wildtype strain with GUS expressed under *pforH* | pRD055 | Hyg^R^ | This work |
| *S. formicae ΔforGF: ΦBT1 GUS pforH* | *forGF* deletion strain with GUS expressed under *pforH* | pRD055 | Hyg^R^ | This work |
| *S. formicaa: ΦBT1 GUS pforZ* | Wildtype strain with GUS expressed under *pforZ* | pRD058 | Hyg^R^ | This work |
| *S. formicae ΔforZ: ΦBT1 GUS pforZ* | *forZ* deletion strain with GUS expressed under *pforZ* | pRD058 | Hyg^R^ | This work |

**Table S3:** ePACs and plasmids used or generated in this study

| **Plasmid** | **Description** | **Resistance** | **Reference** |
| --- | --- | --- | --- |
| pUZ8002 | RK2 derivative with a mutation in *oriT* | Kan^R^ | ^7^ |
| pMS82 | *ori,* pUC18, *hyg, oriT,* RK2, int ΦBT1 | Hyg^R^ | ^8^ |
| pIJ773 | *^-^ aac(3)IV oriT bla* | Apr^R^ | ^9^ |
| pR9604 | pUB307 derivative | Carb^R^ | ^10^ |
| *pESAC-13* 215-G | *aph*II, *tsr* | Kan^R^/Tsr^R^ | BioS&T and ^4^ |
| pCRISPomyces-2 | *AprR, oriT, reppSG5(ts), oriColE1, sSpcas9,* synthetic guide RNA cassette | Apr^R^ | ^11^ |
| pIJ10257 | *oriT*, ΦBT1 *attB-int*, *ermEp**, pMS81 backbone | Hyg^R^ | ^12^ |
| pMF96 | ΦBT1 *attB-int*, uidA CDS, GUS plasmid | Hyg^R^ | ^13^ |
| pIJ10740 (pMF23) | ΦC31 *attB-int, ermEp** | Apr^R^ | ^13^ |
| pRD026 | pCRISPomyces-2 f*orJ* flanking DNA and gRNA | Apr^R^ | This work |
| pRD027 | pCRISPomyces-2 f*orGF* flanking DNA and gRNA | Apr^R^ | This work |
| pRD028 | pCRISPomyces-2 *ForZ* flanking DNA and gRNA | Apr^R^ | This work |
| pRD030 | pMS82 *pforM* forJ | Hyg^R^ | This work |
| pRD031 | pMS82 *pforG* forGF | Hyg^R^ | This work |
| pRD032 | pMS82 *pforZ* forZ | Hyg^R^ | This work |
| pRD034 | pMS82 *pforM* f*orJ* *3x Flag* | Hyg^R^ | This work |
| pRD035 | pMS82 *pforG* f*orGF* *3x Flag* | Hyg^R^ | This work |
| pRD036 | pMS82 *pforZ* *forZ* *3x Flag* | Hyg^R^ | This work |
| pRD037 | *pESAC-13* 215-G *1-4* *aac(3)IV oriT* | Kan^R^/Tsr^R^ | This work |
| pRD038 | *pESAC-13* 215-G *1-7* *aac(3)IV oriT* | Kan^R^/Tsr^R^ | This work |
| pRD039 | *pESAC-13* 215-G *32-43* *aac(3)IV oriT* | Kan^R^/Tsr^R^ | This work |
| pRD040 | *pESAC-13* 215-G *36-43* *aac(3)IV oriT* | Kan^R^/Tsr^R^ | This work |
| pRD054 | pMF96 pforGF GUS | Hyg^R^ | This work |
| pRD055 | pMF96 pforH GUS | Hyg^R^ | This work |
| pRD058 | pMF96 pforZ GUS | Hyg^R^ | This work |
| pRD062 | pMF96 pforJ GUS | Hyg^R^ | This work |
| pRD063 | pMS82 *pforJ* forJ | Hyg^R^ | This work |
| pRD064 | pIJ10257 forJ | Hyg^R^ | This work |

**Table S4:** Primers designed and used for this study (5’-3’). Capital bases indicate overhangs, restriction sites etc.

| **Primer name** | **Description** | **Sequence** |
| --- | --- | --- |
| pCRISP Test F | Test XbaI site pCRISP2 For | aggctagtccgttatcaacttgaaa |
| pCRISP Test R | Test XbaI site pCRISP2 Rev | tcgccacctctgacttgagcgtcga |
| Spacer test | Test BbsI site of pCRISP2 | atacggctgccagataaggc |
| ForJ For1 | Repair template *forJ* KO, left flank | gctcggttgccgccgggcgttttttaTCTAGAggtgtgcgcgaagaacggcc |
| ForJ Rev 1 | Repair template *forJ* KO, left flank | GCTGCTGCGACCAGGCGAGCTCGCcactgacgcggtcgttcccg |
| ForJ For 2 | Repair template *forJ* KO, right flank | GCGAGCTCGCCTGGTCGCAGCAGCtgacgtgcttcgagaccgcc |
| ForJ Rev 2 | Repair template *forJ* KO, right flank | gcaacgcggcctttttacggttcctggccTCTAGAcctcttcatgttcctggtgggcc |
| ForJ gRNA For | sgRNA *forJ* deletion | ACGCtgccgacaccttctccatga |
| ForJ gRNA Rev | sgRNA *forJ* deletion | AAACtcatggagaaggtgtcggca |
| ForJ KO Test 1F | Test *forJ* deletion in genome | cctcttcggtgagcgcttcgagg |
| ForJ KO Test 2R | Test *forJ* deletion in genome | cctgttggacttcgcgcaggc |
| ForJ KO Test 2F | Test *forJ* deletion in genome | gtacgccaggaggacgtgcg |
| ForJ KO Test 1R | Test *forJ* deletion in genome | gccgacgcggcacttctatcc |
| ForZ For1 | Repair template *forZ* KO, left flank | gctcggttgccgccgggcgttttttaTCTAGAcgaacaggccgacgctgaacag |
| ForZ Rev 1 | Repair template *forZ* KO, left flank | GCTGCTGCGACCAGGCGAGCTCGCcatggcttgaagtccagcacgtcc |
| ForZ For 2 | Repair template *forZ* KO, right flank | GCGAGCTCGCCTGGTCGCAGCAGCtcatccgtacctggcagctcgtcg |
| ForZ Rev 2 | Repair template *forZ* KO, right flank | gcaacgcggcctttttacggttcctggccTCTAGAccgaggcggacggatcgcgtcc |
| ForZ gRNA For | sgRNA *forZ* deletion | ACGCgtcggcggtcaactcgactg |
| ForZ gRNA Rev | sgRNA *forZ* deletion | AAACcagtcgagttgaccgccgac |
| ForZ KO Test 1F | Test *forZ* deletion in genome | gccggtgccgaaccggacgc |
| ForZ KO Test 2R | Test *forZ* deletion in genome | cgcacgccgccacgacgagc |
| ForZ KO Test 2F | Test *forZ* deletion in genome | cgcacgccgccacgacgagc |
| ForZ KO Test 1R | Test *forZ* deletion in genome | cgcacgccgccacgacgagc |
| ForGF For1 | Repair template *forGF* KO, left flank | gctcggttgccgccgggcgttttttaTCTAGAggagccggtcttggccatctgc |
| FoGF Rev 1 | Repair template *forGF* KO, left flank | GCTGCTGCGACCAGGCGAGCTCGCggcagcctcgttcacagcagc |
| ForGF For 2 | Repair template *forGF* KO, right flank | GCGAGCTCGCCTGGTCGCAGCAGCtgaggctcaggcgggttcgatgg |
| ForGF Rev 2 | Repair template *forGF* KO, right flank | gcaacgcggcctttttacggttcctggccTCTAGAcgagatcgtcatccacgcgcc |
| ForGF gRNA For | sgRNA *forGF* deletion | ACGCtggcgaagatgttgcgcaga |
| ForGF gRNA Rev | sgRNA *forGF* deletion | AAACtctgcgcaacatcttcgcca |
| ForGF KO Test 1F | Test *forGF* deletion in genome | gcagttcctggacgatgcgc |
| ForGF KO Test 1R | Test *forGF* deletion in genome | cgagggtctggagaacgcgc |
| ForGF KO Test 2F | Test *forGF* deletion in genome | cgtcggcaccttctaccaccg |
| ForGF KO Test 2R | Test *forGF* deletion in genome | gcctgcgtgattcatcggctg |
| pMS82 f*orJ pforM* F1 | *forJ* complementationunder *pforM* | gccgagaaccTAGGATCCAAGCTTgatgccggtgagcagggcgag |
| pMS82 *forJ pforM* R1 | *forJ* complementationunder *pforM* | ggcgccgtggtcgtggtcataccggctcccatcggttgctg |
| pMS82 *forJ pforM F2* | *forJ* complementationunder *pforM* | cagcaaccgatgggagccggtatgaccacgaccacggcgcc |
| pMS82 *forJ pforM R2* | *forJ* complementationunder *pforM* | CTGGTACCATGCATAGATCTAAGCTTgcggaggcggaccgtgcctag |
| pMS82 *forZ* F | *forZ* complementation | gccgagaaccTAGGATCCAAGCTTccggtcaccacccattggag |
| pMS82 *forZ* R | *forZ* complementation | CTGGTACCATGCATAGATCTAAGCTTaggagttgtgcgccctcgc |
| pMS82 *forGF* *F* | *forGF* complementatio | gccgagaaccTAGGATCCAAGCTTcgtgtaccccctgtgcacg |
| pMS82 *forGF* R | *forGF* complementatio | CTGGTACCATGCATAGATCTAAGCTTccgctgctcgccatcgaac |
| pMS82 TEST For | Test HindIII site pMS82 For | gcaacagtgccgttgatcgtgctatg |
| pMS82 TEST Rev | Test HindIII site pMS82 Rev | gccagtggtatttatgtcaacaccgcc |
| ForJ-3xFLAG Rev | *forJ 3x*Flag for ChIP | gcctgaaccgcctccaccgtgccccgcgggcacctg |
| ForJ-3xFLAG For | *forJ 3x*Flag for ChIP | caggtgcccgcggggcacggtggaggcggttcaggc |
| FLAG-pMS82 R | 3*x*Flag in pMS82 for ChIP | CTGGTACCATGCATAGATCTAAGCTTtcaCTTGTCGTCATCGTCCTTG |
| pMS82 ForF prom F | *forGF 3x*Flag for ChIP | gccgagaaccTAGGATCCAAGCTTtcgtgtaccccctgtgcacg |
| ForF-prom R | *forGF 3x*Flag for ChIP | ggtcaccacggtctgcatagcagcctccccggttcg |
| ForGF F | *forGF 3x*Flag for ChIP | cgaaccggggaggctgctatgcagaccgtggtgacc |
| ForF-3xFLAG R | *forGF 3x*Flag for ChIP | gcctgaaccgcctccaccgccccggtcgccctgcg |
| FLAG-pMS82 For | 3*x*Flag in pMS82 for ChIP | cgcagggcgaccggggcggtggaggcggttcaggc |
| ForZ-3xFLAG Rev | *forZ 3x*Flag for ChIP | gcctgaaccgcctccaccccgctcgcacgccgccacg |
| ForZ-3xFLAG For | *forZ 3x*Flag for ChIP | cgtggcggcgtgcgagcggggtggaggcggttcaggc |
| ForJ Test F | To check expression by RT-PCR | gcaaggcggcgcagagcg |
| ForJ Test R | To check expression by RT-PCR | gccgacaccttctccatgagg |
| ForZ Test F | To check expression by RT-PCR | gaaccggacgcagccgcag |
| ForZ Test R | To check expression by RT-PCR | cctcgacgcgtgccacgag |
| ForF Test F | To check expression by RT-PCR | cctcgacgcgtgccacgag |
| ForF Test R | To check expression by RT-PCR | gcgaccagggtcatgacctcg |
| pMF96 HindIII Test For | Test HindIII site of pMF96 | gctcaatcaatcaccggatcc |
| pMF96 HindIII Test Rev | Test HindIII site of pMF96 | catgtccgtacctccgttg |
| 16S r RNA For | qPCR reference gene | cgggtctgcagtcgatacgg |
| 16S r RNA Rev | qPCR reference gene | gctttcgctcctcagcgtcag |
| MLK transcript For | qPCR expression | ctgatcttcggtgccttcctgtcc |
| MLK transcript Rev | qPCR expression | cggcgagcagtccgaggtc |
| J transcript For | qPCR expression | ccgaccgtgcggaaactcg |
| J transcript Rev | qPCR expression | gtccggatccacatgccgc |
| HI transcript For | qPCR expression | ccttcgagttcgtcgtggacg |
| HI transcript Rev | qPCR expression | gctgctcggcgaccagatc |
| GF transcript For | qPCR expression | gctccaccactacgaacagcg |
| GF transcript Rev | qPCR expression | ggagcgagtcctcgatcacg |
| TSRABCDE transcript For | qPCR expression | cgacaccatcgacaccgcc |
| TSRABCDE transcript Rev | qPCR expression | cgttccactccacgaccacc |
| UVWXY transcript For | qPCR expression | gcagcttctccaggagttcc |
| UVWXY transcript Rev | qPCR expression | gccaagaagatcctcgacagg |
| Z transcript For | qPCR expression | ctcatccggctcgtcacgc |
| Z transcript Rev | qPCR expression | cagatggcggttggcgagc |
| AA transcript For | qPCR expression | gaccggaggaacgcctgg |
| AA transcript Rev | qPCR expression | cggtgtcgaggtccttgctc |
| *pforH-G*F F For | *pforG* in pMF96 | AAAAAcatatggcgctgctcacggtcatcg |
| *pforH-G*F Rev | *pforG* in pMF96 | AAAAActcgaggcagcctcgttcacagcag |
| *pforZ-AA* F For | *pforZ* in pMF96 | AAAAAcatatggaatccctgacgcgccgcg |
| *pforZ-AA* F Rev | *pforZ* in pMF96 | AAAAActcgaggacgatggtggtgtcgagcac |
| *forJ* pMF96 360 bp prom for | *pforJ* in pMF96 | caattaatctagaggatccatatgcctgttcgtcgcggtggc |
| *forJ* pMF96 360 bp prom rev | *pforJ* in pMF96 | gtacctccgttgctcgactcgaggttcgctcacctctgctgtgacg |
| *pforGF-H* For | *pforH* in pMF96 | AAAAAcatatggcagcctcgttcacagcagc |
| *pforGF-H* Rev | *pforH* in pMF96 | AAAAActcgaggcgctgctcacggtcatcg |
| pMS82 *forJ pforJ* F1 | *forJ* complementationunder *pforJ* | gccgagaaccTAGGATCCAAGCTTcctgttcgtcgcggtggc |
| pMS82 *forJ pforJ* R1 | *forJ* complementationunder *pforJ* | ggcgccgtggtcgtggtcatgttcgctcacctctgctgtgacg |
| pMS82 *forJ pforJ* F2 | *forJ* complementationunder *pforJ* | cgtcacagcagaggtgagcgaacatgaccacgaccacggcgcc |
| pMS82 *forJ pforJ* R2 | *forJ* complementationunder *pforJ* | CTGGTACCATGCATAGATCTAAGCTTgcggaggcggaccgtgcctag |
| pIJ10257 *forJ* F | *forJ* complementationunder *pErmE** | gtctagaacaggaggccccatatgatgaccacgaccacggcgc |
| pIJ10257 *forJ R* | *forJ* complementationunder *pErmE** | catgagaacctaggatccaagcttggaacgaccgcgtcagtgcc |
